# zFISHer: Automated 3D Registration, Detection, and Colocalization with Interactive Curation for Sequential Multiplexed FISH

**DOI:** 10.64898/2026.05.19.726314

**Authors:** Seth Staller, Virginia Valentine, Steven Burden

**Affiliations:** St. Jude Children’s Research Hospital, Memphis, TN 38105, USA

## Abstract

**Summary:** Sequential multiplexed fluorescence in situ hybridization (FISH) enables spatially resolved molecular profiling in cell monolayers, but analyzing puncta colocalization across three-dimensional (3D) datasets remains a labor-intensive bottleneck. zFISHer is an open-source application built on the napari viewer that provides complete automation of sequential FISH image processing in conjunction with interactive user-curation tools. zFISHer provides end-to-end analysis of paired FISH datasets, encompassing nuclear segmentation, automated puncta detection on unaligned z-stacks, multi-round image registration via translation-constrained RANSAC with optional B-spline deformable warping, precise transformation of puncta coordinates into aligned space, consensus nuclei generation, interactive editing with real-time collision detection, and pairwise and tri-channel colocalization analysis with statistics. This includes a “Fishing Hook” raycasting algorithm that enables users to locate puncta at their true 3D centroids by identifying intensity maxima along the camera ray, eliminating manual z-slice navigation, complemented by a sub-voxel volume optimization. The included batch processing mode enables high-throughput unattended analysis of multiple experimental datasets.

**Availability and Implementation:** zFISHer is open source under the MIT license, freely available on GitHub: https://github.com/stjude/zFISHer. The example dataset (deconvolved ND2 image stacks) is archived on Zenodo at https://doi.org/10.5281/zenodo.20288536. zFISHer is developed in Python utilizing the napari viewer for the interface. Documentation and expected test outputs for the sample dataset are available on the GitHub: https://github.com/stjude/zFISHer. To report an issue using zFISHer or contributing to it, please file an issue in the GitHub repository: https://github.com/stjude/zFISHer/issues.

**Supplementary Information:** Supplementary data are available online.

## 1 INTRODUCTION

Sequential multiplexed fluorescence in situ hybridization (FISH) has become a widely adopted technique for spatially mapping molecular targets within cells (Lubeck et al., 2014; Chen et al., 2015; Eng et al., 2019). In these experiments, cells undergo multiple rounds of labeling and imaging to produce volumetric image stacks that must be properly registered and analyzed together to accurately identify colocalized signals. This enables characterization of localization patterns of subcellular target distributions in disease-relevant cell populations.

Due to multiple rounds of processing and acquisition and the required accuracy for distances at the molecular scale, the ideal analysis of sequential FISH data faces three intertwined challenges. Cross-round acquisition drift and nonlinear elastic deformation must be aligned to a shared coordinate space before volumes can be meaningfully compared. Puncta annotation must operate on z-stacks rather than 2D maximum intensity projections, because it is typical for puncta in crowded fields to share XY positions but exist at different depths in capture volumes. Manual puncta curation is time-consuming, as multiple rounds of capture contain many z-slices that must be scrolled through individually, and a cursor on a 2D screen has no meaningful depth for annotation in 3D volumes. Existing tools will address one or two of these challenges in isolation, which necessitates programming expertise to stitch workflows across multiple codebases. Those contemporary tools that do operate in 3D typically lack interactive puncta curation or require dedicated environments that are suboptimal for the open, Python-native ecosystem of modern workflows.

Starfish (Axelrod et al., 2021) provides programmatic pipelines for image-based transcriptomics and allows outputs to be viewed in napari but requires programming skills to develop an analysis flow. Punctatools (Baggett et al. 2022), RS-FISH (Bahry et al., 2022), and Big-FISH/FISH-quant v2 (Imbert et al., 2022) provide automated 3D spot detection, with Punctatools and Big-FISH supporting pairwise colocalization, but lack multi-round registration or interactive tools for editing. The most robust open source tool, pyHiM (Devos et al, 2024), is an automated python pipeline for multiplexed DNA-FISH and included multi-round registration and spot detection, but lacks interactive curation and is designed for chromatin tracing. No existing, open source, pythonic tool integrates the three capabilities of automated detection, 3D registration, and interactive curation, into a single programmatic-free environment.

To address these gaps, zFISHer is an open-source application utilizing the napari viewer (Ahlers et al., 2023) that provides a fully automatable toolset for segmentation, registration, detection, curation, and colocalization analysis of multi-round sequential 3D FISH data within a single unified environment. zFISHer analysis automation executes per-round nuclear segmentation (modified classical watershed or Cellpose; Stringer et al., 2021), two-stage rigid then deformable registration, 3D puncta detection with four algorithm options (local maxima, LoG, DoG, modified radial symmetry), and pairwise and tri-channel colocalization. zFISHer architecture was deliberately designed to follow a detect-then-transform approach to maximize accuracy and avoid interpolation-derived artifacts. Additionally, zFISHer also includes a suite of interactive refinement tools, including a nuclei mask editor, manual puncta picking, and a Fishing Hook ray-casting algorithm that identifies true 3D puncta centroids from any viewing angle with intensity-based depth calculation. The batch analysis mode, driven by a configurable parameter spreadsheet, enables unattended processing of multiple, large datasets. All modules are architecturally distinct from the interface, allowing scriptable pipeline development, and a headless command-line interface. A comparison of zFISHer’s capabilities to published tools is provided in Supplementary Table S1.

## 2 IMPLEMENTATION

zFISHer is developed in Python and utilizes the napari viewer through a magicgui widget framework. The architecture separates core algorithms from the graphical interface and thus enables both headless automation and interactive operation. The analysis is organized into five sequential steps: (1) session and I/O, (2) nuclei segmentation, (3) puncta detection on unaligned z-stacks, (4) cross-round alignment and consensus, (5) colocalization and analysis. Each zFISHer analysis is persisted as a reloadable session file with linked reports and logging for provenance, allowing users to save, reload, refine, and share analyses. The zFISHer analysis workflow is diagrammed in Supplementary Figure S2. A more detailed explanation of the implementation is given in Supplementary Methods S1.

### 2.1 Nuclei segmentation and consensus

Per-nucleus colocalization of puncta requires accurate 3D segmentation to properly map puncta to their proper nuclei. zFISHer independently segments on the unregistered nuclear channels prior to registration to allow flexibility during manual puncta curation and inform downstream registration and warping. zFISHer provides a modified classical watershed pipeline consisting of downsampling, Gaussian smoothing, Otsu thresholding, anisotropy-corrected distance-transform peak finding, and over-segmentation merging (Otsu, 1979. Additionally, native integration of Cellpose is included for challenging fields (Stringer et al., 2021). Each segmentation mask is annotated with a unique ID and a centroid for downstream processing, such as puncta assignment (Section 2.2) and consensus nuclei mask generation following downstream cross-round z-stack alignment (Section 2.4). The interactive mask editor permits manual mask refinement, including painting, deletion, and z-dimensional extrusion. While the underlying segmentation algorithms (Otsu thresholding, watershed, Cellpose) are well-established, zFISHer’s contribution lies in anisotropy-aware modifications, over-segmentation merging, per-round mask editing, and downstream cross-round consensus generation in a unified environment.

### 2.2 Detect-then-transform puncta processing

A core design decision of zFISHer was to perform puncta detection on unaligned image inputs and then transform the coordinates into the unified canvas space. This approach allows detection on the original signal to avoid interpolation artifacts. Four 3D detection algorithms are included: local maxima, Laplacian of Gaussian (LoG), Difference of Gaussian (DoG), and a modified radial symmetry algorithm (Parthasarathy, 2012). The radial symmetry implementation was modified for 3D detection, and refines each candidate to sub-voxel precision via weighted least-squares gradient-ray intersection. LoG, DoG, and radial symmetry utilize anisotropic Gaussian kernels with sigma vectors (σ*dz/dx,σ,σ) to match the elongated geometry typical in FISH z-stacks. Pre-processing background subtraction is provided via a per-slice white top-hat filter. Each detected puncta is annotated with coordinates, intensity, SNR, and a unique ID.

### 2.3 The Fishing Hook algorithm

Manual annotation of puncta in 3D FISH volumes is a time-consuming step of the analysis. A common concession is to reduce the image volume to 2D maximum intensity projections. However, this fails to resolve puncta at similar XY positions but different Z depths typical in crowded fields. Navigating individual z-slices is slow, subjective, and imprecise, as puncta in FISH fields tend to span across adjacent slices, making the identification of the true signal centroid challenging, if not impossible. Automated detection is imperfect and requires manual verification in practice.

Despite advances in 3D image processing and visualization, resolving 3D depth remains a challenge for 2D interactions because cursors do not have meaningful depth. Intensity-based depth resolution was popularized by Vaa3D (Peng et al, 2014) for tracing neurons in standalone C++ environments. It is available in commercial tools such as Imaris (Bitplane), but the functionality is lacking in modern, open-source, Python-native ecosystems for FISH analysis workflows. Contemporary Python-based tools such as napari-threedee (Yamauchi and Burt, 2023) focus on plane-based interactions and geometric intersections but do not leverage image data for coordinate snapping.

The Fishing Hook algorithm was developed for zFISHer to provide a fast and intuitive method for users to identify unannotated puncta directly in the 3D image volume. The algorithm uses the camera orientation and cursor position to construct a multi-point ray through the volume that is normalized via anisotropy-aware voxel scaling. Intensities are sampled at each point position along the ray to identify the brightest voxel candidate. Then, an optional local 3D neighborhood search refines the candidate position to the highest intensity peak in a defined radius to correct for cursor offset. Redundant annotations are handled by an integrated collision detection threshold. The implementation is detailed in Supplementary Methods 1.6.

### 2.4 Multi-round registration and warping

Multiple rounds of sample processing between rounds of image acquisition produces inter-round drift. Global transformations can correct the rigid components, but elastic nonlinear deformation presents the greatest challenge for image alignment for accurate molecular spatial profiling. zFISHer addresses inter-round volume alignment with a three-phase process: RANSAC-based rigid alignment on nuclei centroids (Fischler and Bolles, 1981), channel-type-aware deformation for refinement, and coordinate-space unification of nuclei masks and puncta.

Rigid alignment uses R1 and R2 nuclei centroid clouds to estimate a coarse shift by vector voting on all pairwise differences. Nearest-neighbor correspondences are identified with a 3D KD-tree. A custom 3-DOF translation model is then fit with RANSAC. A translation-only model was chosen to prevent overfitting characteristic of affine models fit to sparse correspondences, and excessive deviation triggers reversion to the coarse shift. Each round volume is then padded to a unified canvas space. The optional second-stage channel deformation applies B-spline transform (Rueckert et al., 1999) to the rigidly aligned R2 channel using 4-node mesh per axis, Mattes mutual information, and L-BFGS-B optimization. The transformation is applied to all R2 image channels with channel-type-aware interpolation: B-spline for nuclear channel and linear for the remaining image channels.

Following alignment and warping, the per-round nuclei masks are unified to a consensus mask via KD-tree with an adaptive distance threshold. Masks can be merged in union or intersection, and nuclei centroids are recomputed upon completion. Puncta coordinates are transformed by applying the canvas offset, rigid shift, and an iterative inverse B-spline, and then re-indexed against the consensus nuclei masks. The implementation is detailed in Supplementary Methods 1.7-1.9.

### 2.5 Colocalization and analysis

Sequential FISH experiments are primarily performed to map molecular spatial profiles of multiple elements within the nucleus. Colocalization analysis in zFISHer is performed on puncta coordinates following registration and warping to ensure accurate inter-round calculation in unified nuclear space. To maintain consistency across datasets, resolutions, and acquisitions, all distance calculations operate in physical units (micrometers) using voxel scaling derived from the input dataset OME-XML metadata. Both pairwise and tri-channel colocalization modes are provided. Pairwise colocalization performs a bidirectional 3D radius search between two channels using a KD-tree. Every unique spot pair within a user-defined threshold is reported as a positive hit. Tri-channel colocalization uses a greedy assignment strategy on an anchor channel and two additional channels to mimic a biological model of multiple elements localizing to a single spatial locus. While more rigorous approaches for tri-channel colocalization exist, such as MultiMatch (Naas et al., 2024), the greedy strategy prioritizes interpretability and speed. All puncta of both child channels within the user-defined radius threshold of an anchor channel puncta are enumerated. The greedy algorithm iteratively picks pairs of channel A/B puncta that are closest to the anchor and removes them from the pool to prevent double counting until no valid triplets remain. For convenience, a separate nearest-neighbor distance list is compiled for every point in every channel against every other channel to produce a comprehensive background distribution. Metrics are compiled for reporting, including per-hit tables, per-nucleus aggregates, and summary statistics. The implementation is detailed in Supplementary Methods 1.10-1.11.

### 2.6 Batch and headless processing

Sequential FISH experiments generate multiple sets of image pairs per study, and manual analysis of each dataset through interactive GUIs is time-consuming and variable across multiple operators. zFISHer addresses this by enabling fully automated execution of all stages of analysis independent of user interaction. The headless architecture can be executed through custom scripting or CLI commands for implementation into larger workflow pipelines. For users without programming experience, a GUI-initiated batch processing mode provides the same automated execution. The multi-sheet input template allows full parameter configuration and definitions for all stages of the analysis. The templates are validated against each input dataset before execution to prevent mid-run failure. All outputs (masks, puncta CSVs, reports, session, logs) from each dataset pair are written to unique subdirectories under a batch parent output root directory for provenance. This enables overnight, unattended, and unbiased processing for large FISH datasets. The finished sessions can then be loaded into zFISHer for refinement, curation, and additional analysis if needed. The implementation is detailed in Supplementary Methods 1.1.

## 3 APPLICATION

zFISHer addresses two primary bottlenecks of sequential FISH analysis: the inter-operator variability and time-intensive nature of navigating individual z-slices to confirm puncta depth. Additionally, the software’s puncta-based processing is signal agnostic and thus applicable to mixed-modality workflows. To demonstrate this, the sample data set combines DNA FISH (HOXB locus, FITC) with protein immunofluorescence (XPO1, Cy5) in human acute myeloid leukemia Molm13 cell monolayers (Supplementary Methods S2; Staller et al., 2026). The analysis performed with zFISHer (Supplementary Methods S3) used automated processing for nuclear segmentation (Figure 1A), puncta detection on unaligned images, RANSAC-based registration with B-spline warping (Figure 1B), and consensus nuclei generation as a single operation across both z-stacks, which contained 1,280 Cy5 puncta and 112 FITC puncta across 55 nuclei (Figure 1C). Compared to the existing manual workflow that took over four hours of manual registration and puncta validation, zFISHer reduced total analysis time to under one hour. The Fishing Hook algorithm was used for curation in the FITC channel, which eliminated the time-consuming need to navigate individual z-slices manually (Figure 1D). The results show an expected signal distribution for per-nucleus frequency of each probe, with the HOXB DNA puncta averaging two per cell characteristic of a diploid nucleus, and the XPO1 showing a pattern consistent with variable protein expression across cells (Figure 1E). The pairwise colocalization results identified 8 of n=112 puncta falling below a 1 μm colocalization threshold, consistent with the expected absence of spatial association between the chromatin locus and nuclear export protein (Figure 1F). This demonstrates zFISHer’s ability to quantify cross-round, cross-channel puncta spatial relationships without bias.

**Figure 1.**
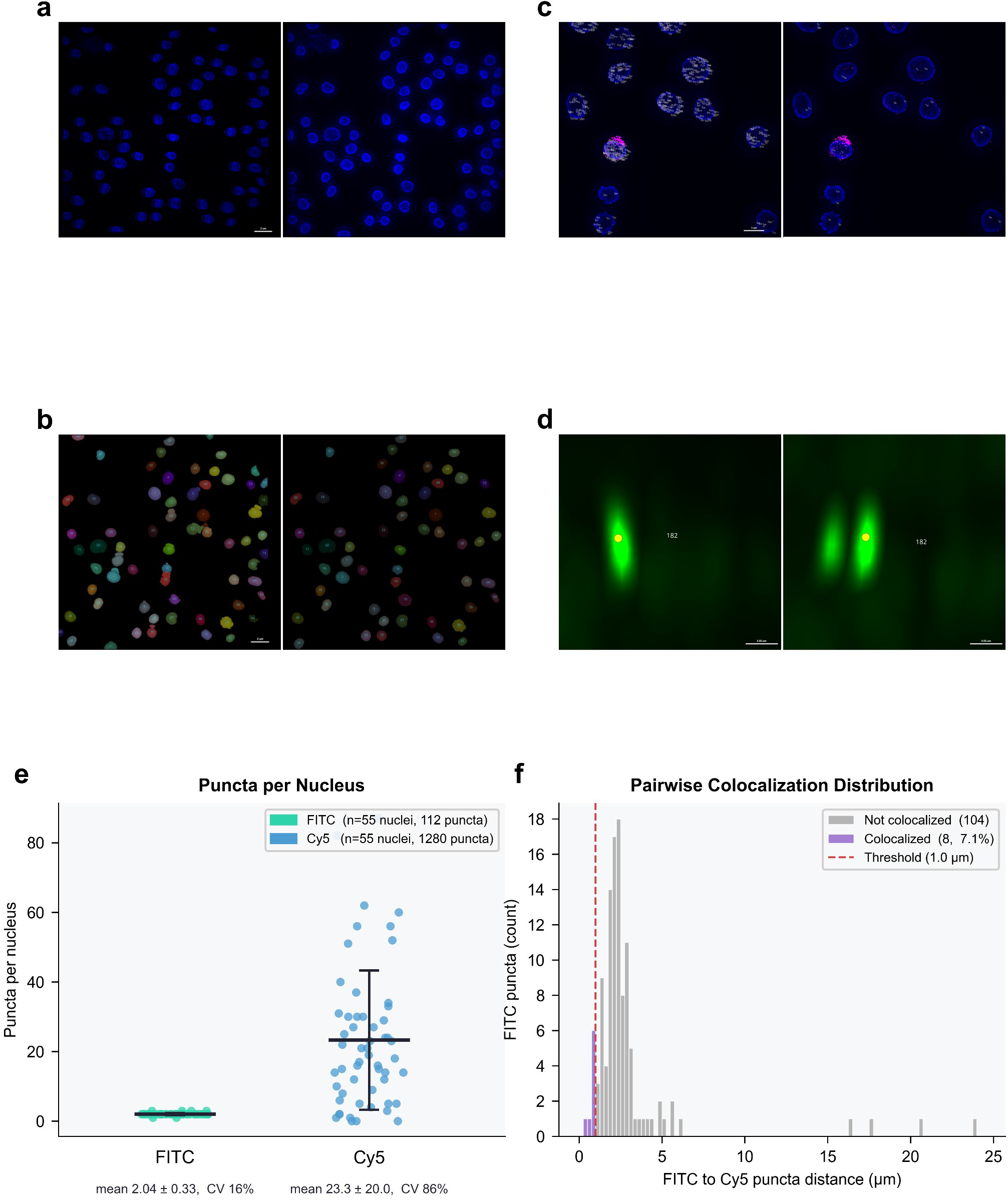
zFISHer enables end-to-end 3D puncta curation and colocalization in sequential multiplexed FISH. a) Overlay of DAPI images from file 1 and file 2 before (left) and after (right) alignment and registration. b) Global nuclear mask with assigned IDs before (left) and after (right) manual cleanup. Cleanup included removal of partial nuclei at image margins and elimination of non-nuclear masks. c) Warped Cy5 channel overlaid with aligned FITC channel. Left: Cy5 with puncta IDs and FITC without puncta IDs. Right: Cy5 without puncta IDs and FITC with puncta IDs. d) Demonstration of Fishing Hook puncta curation in the FITC channel. Left: the algorithm defined by cursor position in viewer identifies and places the punctum (yellow dot) at the axial intensity maximum of the signal, correcting for depth misassignment by volume-based optimization. Right: side view of two axially overlapping puncta. The algorithm correctly selects the rearmost punctum because it corresponds to the brighter intensity maximum, illustrating the algorithm’s ability to disambiguate signals along the optical axis. e) Per-nucleus puncta counts (n=55). FITC puncta (HOXB DNA locus; mean 2.04 ± 0.33, CV 16%) is consistent with two puncta per diploid nucleus. Cy5 (XPO1 protein; mean 23.3 ± 20.0, CV 86%) is consistent with variable expression across cells. This demonstrates zFISHer’s ability to handle mixed-modality datasets. f) Distribution of nearest-neighbor distances of each FITC punctum to its closest Cy5 punctum (n=112). FITC puncta with a Cy5 neighbor falling within the 1.0 µm colocalization threshold (n=8) are drawn in purple. The remaining non-colocalized FITC puncta (n=104) are shown in gray. This demonstrates zFISHer’s ability to resolve between colocalization events (≤ 1.0 μm) and non-colocalized neighbors (> 1.0 μm).

## AVAILABILITY AND REQUIREMENTS

**Project name:** zFISHer

**Project home page:** https://github.com/stjude/zFISHer

**Example dataset (inputs):** https://doi.org/10.5281/zenodo.20288536

**Operating system(s):** Platform independent (Windows, macOS, Linux)

**Programming language:** Python 3.10

**Other requirements:** napari, magicgui, numpy, scipy, scikit-image, SimpleITK, pandas, tifffile, nd2, PySide6, cellpose, openpyxl, qtpy, markdown, packaging, Pillow

**License:** MIT

## ACKNOWLEDGEMENTS

Thank you to Melvin Thomas and the Jeffery Klco Lab for providing the sample that generated the example data images.

