## Supplementary Information for "zFISHer: Automated 3D Registration, Detection, and Colocalization with Interactive Curation for Sequential Multiplexed FISH"

#### **Supplementary Table S1.** Feature comparison of zFISHer with existing tools for FISH image analysis.

| Feature | starfish | Big-FISH / FISH-quant v2 | RS-FISH | punctatools | Quant Punc | napari-threedee | pyHiM | zFISHer |
|---|---|---|---|---|---|---|---|---|
| Native 3D operation | ✓ | ✓ | ✓ | ✓ | – | ✓ | ✓ | ✓ |
| Multi-round image registration | – | – | – | – | – | – | ✓ | ✓ |
| Interactive GUI | – | ~ | ✓ | – | ✓ | ✓ | ~ | ✓ |
| Interactive puncta curation / editing | – | – | – | – | ~ | ~ | – | ✓ |
| Intensity-guided 3D centroid snapping | – | – | – | – | – | – | – | ✓ |
| Automated spot detection | ✓ | ✓ | ✓ | ✓ | ✓ | – | ✓ | ✓ |
| Nuclear segmentation | ~ | ✓ | – | ✓ | ✓ | – | – | ✓ |
| Consensus nuclei across rounds | – | – | – | – | – | – | – | ✓ |
| Pairwise colocalization analysis | – | ✓ | – | ✓ | ✓ | – | ✓ | ✓ |
| Tri-channel colocalization | – | – | – | – | – | – | – | ✓ |
| Detect-then-transform workflow | – | – | – | – | – | – | – | ✓ |
| Per-nucleus statistics | ~ | ✓ | – | ✓ | ✓ | – | – | ✓ |
| Batch processing | ✓ | ✓ | ✓ | ✓ | – | – | ✓ | ✓ |
| napari integration | – | – | – | – | ✓ | ✓ | – | ✓ |

**Supplementary Table S1. Feature comparison of zFISHer with existing tools for FISH image analysis.** Support is indicated (✓ = supported; ~ = partial support; – = not supported). Description of partial support (~) is as follows. Interactive GUI: starfish is code-only; Big-FISH has ImJoy web plugins (~); pyHiM contains parameter configuration GUI but no interactive analysis environment; Puncta curation: QuantPunc allows manual 2D editing (~); napari-threedee enables plane-based 3D annotation (~) but not intensity-guided placement. Nuclear segmentation: starfish provides segmentation on maximum projections (~). Per-nucleus stats: starfish provides cell-by-gene matrices

(~); pyHiM provides per-cell chromatin trace statistics (~). Support was determined from referenced sources documentation (starfish: Axelrod et al. (2021); Big-FISH/FISH-quant v2: Imbert et al. (2022); RS-FISH: Bahry et al. (2022); punctatools: Baggett et al. (2022); QuantPunc: napari-hub; napari-threedee: Yamauchi and Burt (2023); pyHiM: Devos et al. (2024).

### **Supplementary Figure S2.** Overview of the zFISHer workflow.

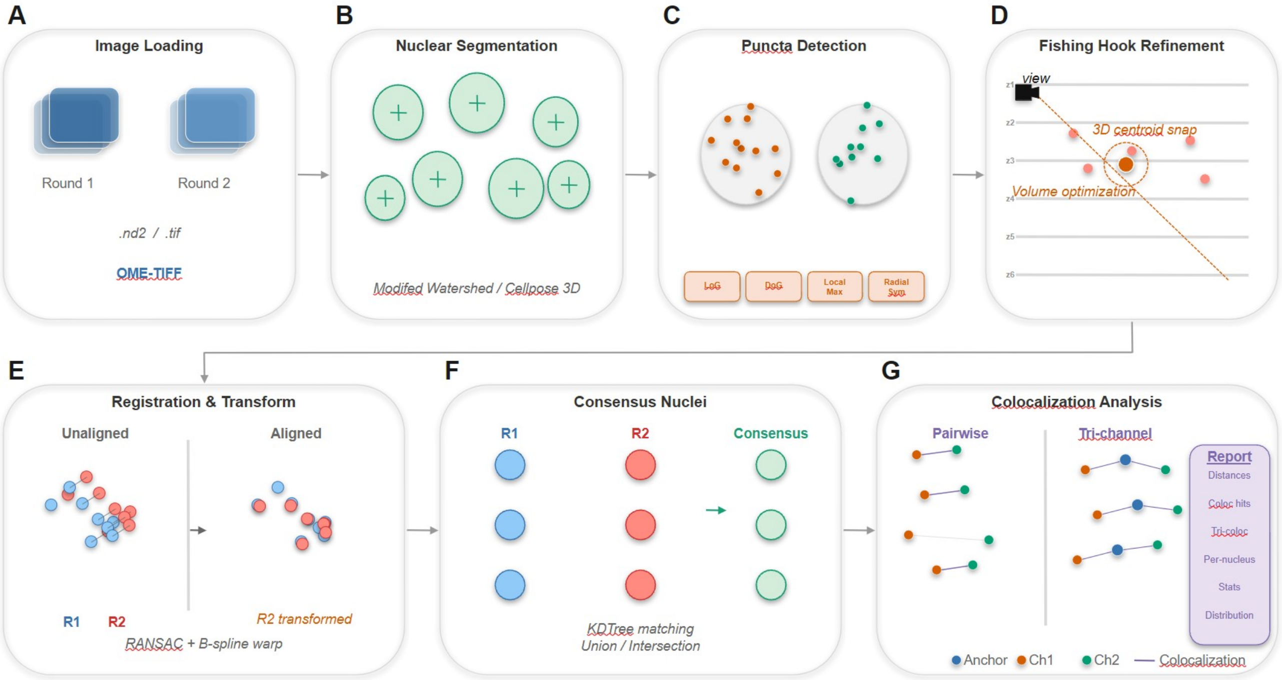


**Figure S2. Overview of the zFISHer workflow.** (A) Sequential FISH images from two rounds are loaded and converted to OME-TIFF standard format. (B) Per-round nuclei channel segmentation and centroid annotation using the modified watershed or Cellpose algorithm. (C) Automated 3D puncta detection on raw, unaligned FISH channels using per-round nuclei masks, using one of the four supplied algorithms: Laplacian of Gaussian (LoG), Difference of Gaussian (DoG), Local Maxima, or modified Radial Symmetry (Radial Sym). (D) Interactive 3D puncta curation using the Fishing Hook ray-casting algorithm with volume optimization. (E) Two-stage registration (translation-constrained vector-voting + RANSAC alignment followed by optional B-spline deformable warping) produces the aligned canvas. Existing puncta coordinates are mathematically transformed into aligned space. (F) Consensus nuclei merging across rounds with extranuclear puncta removal. (G) Pairwise and tri-channel colocalization analysis with per-nucleus statistics and report generation.

### **Supplementary Methods 1.** zFISHer Implementation Details.

#### 1.1 Session, I/O, and batch processing

**Session creation and state.** A new zFISHer session is initialized by specifying an output directory and two input image files corresponding to Round 1 (R1) and Round 2 (R2) of sequential hybridization. On initialization, the session constructs a standardized subdirectory layout (input/, segmentation/, aligned/, reports/, captures/, logs/) within the user-specified output directory and writes a JSON session file (e.g.: zfisher_session_1.json) that persists all the analysis state. The session tracks the output path, input image paths, registration shift vector, processed file registry, colocalization rule sets (pairwise and tri-channel), and per-channel puncta detection parameters. All session writes are thread-safe to support concurrent UI and background-worker access.

**Input formats.** *z*FISHer natively accepts the standardized OME-TIFF and Nikon ND2 files as inputs. Volumetric (3D) z-stacks are required, and 2D images and maximum-intensity projections (MIPs) are rejected with error at load time. ND2 files are immediately auto-converted to OME-TIFF on session initialization using the nd2 library, with voxel spacing, channel names, and acquisition metadata preserved in the OME-XML schema. TIFF files are parsed through tifffile, with a fallback to ImageJ metadata when OME-XML is absent. Each image is represented as an internal FISHSession dataclass containing the raw (Z, C, Y, X) array, voxel dimensions in micrometers (μm), channel labels, and the original metadata dictionary.

**Session persistence and reload.** The session state is serialized to JSON, including file paths, segmentation outputs, puncta coordinate tables, rule definitions, and per-layer processing parameters. Loading an existing session preserves the original session file and creates a new, incremented session file (e.g. “zfisher_session_2.json”) in the output directory to maintain an audit history and prevent accidental overwrites. This allows users to branch analysis, share session files with collaborators, or resume interrupted work without re-running upstream steps.

**Output directory structure.** The output directory of an analysis contains:

- input/ - converted OME-TIFF files, OME-XML metadata dumps, and raw ND2 metadata JSON (if applicable)
- segmentation/ - nuclei masks and centroid arrays for each round
- aligned/ - registration-warped channels and consensus masks in aligned canvas space
- reports/ - puncta coordinate tables (CSV) and analysis reports (XLSX)
- captures/ - user-defined screenshots
- logs/ - per-session log file with all parameters, warnings, and processing timings

**Batch processing.** Batch mode is driven by a multi-sheet input template that is generated directly from the UI. The template includes Instructions, Datasets, Puncta, and Colocalization sheets. The Datasets sheet defines one row per sample, specifying R1/R2 paths, nuclei segmentation method, and channel mappings. The Puncta sheet allows per-channel fine-tuning of algorithmic detection parameters. The Colocalization sheet allows definition of pairwise and tri-channel rules to be applied to sets of channels. Once loaded, a batch template is parsed and validated. Each dataset is then run sequentially through the full pipeline- segmentation, puncta detection, registration, consensus generation, and colocalization analysis – with reports placed in per-dataset subfolders under a common batch output directory. This allows overnight unattended processing of many datasets with consistent parameters.

**Headless operation.** The core analysis modules (zfisher.core.*) are architecturally separated from the napari UI layer. Session initialization, image loading, segmentation, registration, and report generation are all callable without instantiating a viewer, enabling scripted pipelines, automated testing, and integration into larger workflows. The same functions are invoked by the interactive widgets in the napari GUI and the batch processor.

#### 1.2 Nuclear segmentation

Nuclei are segmented independently on each round's user-specified nuclear channel before any inter-round alignment, ensuring that registration operates on landmarks derived from the unaltered signals within the input files. zFISHer natively provides two complementary segmentation backends, selectable per dataset: a modified, classical watershed pipeline and Cellpose.

***Modified classical watershed.*** The default classical pipeline is designed for speed and cleanly-separated nuclei typical of well-spread cell monolayers. The classical pipeline is implemented on top of scikit-image (gaussian, threshold_otsu, watershed, peak_local_max) and SciPy (distance_transform_edt, ndimage). The algorithm executes as follows:

1. *Downsampling.* The input volume is downsampled in Z (every 2$^{nd}$ slice) and XY (by a factor of 0.5) to accelerate subsequent steps without compromising nucleus-level accuracy.
2. *Gaussian smoothing.* A Gaussian filter ($\sigma$= 3 voxels) suppresses high-frequency noise that could fragment thresholded regions.
3. *Otsu thresholding.* An automatic intensity threshold is computed via Otsu's method (Otsu, 1979), producing a binary foreground mask. Connected components smaller than 50 voxels are removed as noise.
4. *Anisotropy-corrected distance transform.* A Euclidean distance transform is computed on the binary mask using the physical voxel spacing (dz, dy, dx) as the sampling vector, so that the distance metric reflects true micrometer distances rather than voxel counts. This prevents over-splitting along the axially elongated Z direction typical of FISH imaging.
5. *Peak detection.* Candidate nucleus centers are identified by local maxima in the distance-transformed volume, separated by at least 7 voxels.
6. Watershed. The candidate points seed a watershed segmentation on the negated distance transform, producing labeled regions constrained to the Otsu foreground mask.
7. *Over-segmentation merging.* A post-processing step finds all pairs of spatially adjacent labels and computes their shared boundary surface as a fraction of the smaller label's total surface. Pairs with a shared boundary exceeding 30% of the smaller label's surface are merged via union-find, on the assumption that such thin interfaces reflect a single nucleus split by the watershed rather than two distinct nuclei in contact. This step resolves the common failure mode of distance-transform watershed on densely packed nuclei.
8. *Rescaling.* Masks are resized back to the original image shape with nearest-neighbor interpolation, and centroids are extracted from the final labeled volume.

**Cellpose Integration.** zFISHer provides native integration with Cellpose (Stringer et al, 2021), intended for datasets with densely packed, morphologically complex, or touching nuclei. The input volume is downsampled (Z every 2$^{nd}$ slice, XY by 0.25) and fed to Cellpose's pretrained nuclei model in 2D-per-slice mode with cross-slice label stitching (IoU-based stitching threshold of 0.3). Cell diameter is auto-estimated by Cellpose's built-in size model, which predicts the average nucleus diameter from the image and rescales accordingly before segmentation. GPU acceleration is used automatically when available, otherwise the model runs on CPU. The resulting volume is resized back to the input shape with nearest-neighbor interpolation. The same over-segmentation merging step used in the classical pipeline is optionally applied to the Cellpose output.

**Segmentation output.** Both backends return a (Z,Y,X) labeled integer mask and an (N, 3) array of nucleus centroids in the original image coordinate system. Masks are saved as TIFF files in the segmentation/ subdirectory. Centroids are saved as NumPy arrays for use in downstream registration and consensus generation.

#### 1.3 Interactive mask editing

Algorithmic nuclei segmentation unavoidably produces artifacts, such as false positives near the z-stack edges, under-segmentation of nearby nuclei, over-segmentation into multiple masks, and missed nuclei. zFISHer provides a mask editor equipped with a set of topological and geometric operations for correcting errors directly in the napari viewer canvas space.

**Editing operations.** The mask editor supports the following actions:

- *Paint and erase.* Draw new or erase existing nuclei masks with a modified configuration of napari's native brush tools. An ID can be specified to add voxels to an existing mask. New masks can be assigned a new integer ID based on the existing array.
- *Merge.* Two masks can be combined into a single ID.
- Extrude. A mask can be filled through its entire z axis by projecting the union of it XY footprints across slices. The fill respects existing voxel labels.
- *Delete.* A specified ID can be removed with its voxels set to background. Additionally, the entire mask layer can be cleared for fresh segmentation or manual painting.
- *Hover-delete mode.* A toggled mode highlights a nuclei mask in red under the cursor via a custom napari CyclicLabelColormap expansion that assigns each label a unique color slot. A hotkey press removes the highlighted mask and ID, allowing for rapid removal.

**Undo history.** A diff-based undo stack (_MaskUndoStack) records the changed voxel indices alongside their previous values, rather than full array copies. Each editing operation begins with a snapshot and ends with a diff computation against the snapshot. This bounds memory usage, even on large volumes.

**Saving and ID synchronization.** Editing triggers autosave of the modified mask back to its TIFF on disk, and ID centroid and numeric label overlays are recomputed for subsequent operation reference.

**Downstream integration.** The editor writes to the same mask file used for registration, consensus generation, and puncta assignment. Corrections automatically propagate through the entire pipeline without re-import.

#### 1.4 Automated puncta detection

Puncta detection operates on unaligned and unwarped images using per-round nuclei masks for nuclear assignment. Detection before registration ensures spot identification operates on the original signal to avoid interpolation artifacts from B-spline transformation.

**Detection algorithms.** zFISHer includes four 3D spot-detection algorithms:

- *Local maxima.* Fast, intensity-based peak identification utilizing skimage.feature.peak_local_max. There is optional prefiltering with a 3D Gaussian for noise reduction, and a configurable minimum separation distance between peaks. It is ideal for well-separated and high contrast puncta.
- *Laplacian of Gaussian (LoG).* Scale-space blob detection via skimage.feature.blob_log. The Gaussian kernel uses anisotropic sigma vector ($\sigma \cdot z\_scale$, $\sigma$, $\sigma$) where z_scale = dz/dz to match the kernel aspect ratio with the voxel geometry. Well-suited for similarly sized puncta blobs with higher background noise.

- *Difference of Gaussian (DoG).* A fast approximation of LoG via skimage.feature.blob_dog. It also utilizes the anisotropic sigma vector. This is ideal for faster run times on larger dataset volumes with puncta that are well-detected with LoG and can tolerate loss of accuracy.
- *Radial symmetry.* The slowest but most precise algorithm for sub-voxel localization. It is based on Parthasarathy's radial symmetry transform (Parthasarathy, 2012) but is extended to 3D. Candidate peaks are identified by local maxima, then each peak is refined by computing local voxel intensity gradients that are then solved by a weighted least-squares system for the point of convergence of the gradient rays. The approach is used when maximum accuracy is necessary.

**Preprocessing.** A per-slice white top-hat filter is available to suppress low-frequency background gradients. It is disabled by default, due to its tendency to attenuate dim puncta.

**Metrics.** Each puncta is annotated with two per-spot metrics that are retrained with coordinates for downstream filtering:
- *Peak intensity.* The image value at the spot's center voxel.
- *Signal-to-noise ratio (SNR).* The ratio of peak intensity to the median intensity in a local 2D neighborhood on the spot's z-slice. A 3D neighborhood is not used because FISH puncta typically span less than two z-slices, and sampling local background would artificially depress the local neighborhood intensity and inflate SNR.

**Nuclear assignment.** Each detected spot is assigned to a nucleus ID by indexing the per-round mask at its ZYX coordinates. Spots falling outside mask volumes receive ID 0. The discard of extranuclear puncta can be toggled to restrict analysis to signal within segmented nuclei via the nuclei_only parameter.

**Output.** Puncta are saved as per-channel CSV files with the schema (Z,Y,X, Nucleus_ID, Intensity, SNR) in the reports/ subdirectory.

#### 1.5 Manual puncta curation

Algorithmic spot detection produces artifacts and errors such as missed dim puncta, overlapping puncta detected as a single event, and false positives from debris or out-of-focus elements near the edge of the volume. Similar to the mask editor, zFISHer offers a dedicated puncta editor widget for curation and refinement of channel puncta directly in the napari canvas view space.

**Editing operations.** The puncta editor supports the following actions:
- *Add.* Puncta can be added on each channel in 2D (per slice) at a given ZYX position.
- *Fishing Hook (see 1.6).* This novel ray-casing placement algorithm identifies puncta in the 3D volume canvas space. This tool is ideal for 3D curation.
- *Move.* Selected puncta can be repositioned via drag.
- *Delete.* Selected puncta can be deleted in bulk, and a keyboard shortcut allows rapid deletion of puncta under the cursor. An entire channel layer can be cleared of puncta and their IDs.

**Undo history.** A purpose-based undo stack (_PunctaUndoStack) records data and layer snapshots. Data snapshots are taken of a layer's coordinate array and feature table prior to each add/delete/move operation. Before layer deletion, the state (data, features, scale, translate, point size, face color, text configuration, blending) is captured. The stack has a fixed depth and is cleared on layer switch.

**Saving and ID synchronization.** Each layer carries a pandas feature table (nucleus ID, intensity, SNR, and Source) inherited from automatic detection. Provenance is preserved, allowing the tracking of the puncta origin (algorithm vs. manual). Editing triggers autosave of the layer's combined coordinates and feature table back to the CSV in the reports/ subdirectory. Puncta IDs are recomputed for subsequent operation reference.

**Downstream integration.** The source CSV file in the reports/ subdirectory is utilized by downstream coordinate transformation, consensus nuclei ID reassignment, and colocalization analysis. Corrections automatically propagate through the entire pipeline without re-import.

#### 1.6 The Fishing Hook algorithm

Manual curation of FISH puncta in 3D volumes is bottlenecked by navigation through individual z-slices to identify, verify, and annotate their true signal coordinate position. A 3D projection or rendering allows easy visualization for identification but does not solve the annotation issue, because a cursor has no meaningful depth in an image volume displayed on a 2D screen. zFISHer's Fishing Hook algorithm solves this annotation issue by allowing the user to identify a punctum from any camera angle and automatically snapping it to the true 3D position of the underlying signal in the source. The algorithm relies only on raw image data and camera orientation and thus is robust to atypical shapes or signals that might confuse a trained and learned detector.

**Algorithm process.** Upon holding the Fishing Hook hotkey and clicking on the canvas, the click position is converted into a world-space coordinate on the viewing plane. From this seed position, the algorithm proceeds as follows:

1. *Camera orientation.* The current camera view position is read from the napari viewer (viewer.camera.view_direction) in world coordinates. The world-space direction vector is generated from the combination of the camera orientation and cursor position.
2. *Anisotropy correction.* The world-space direction vector is converted to voxel space by dividing the layer scale voxel spacing. The converted vector is normalized to unit length for geometrically consistent stepping.
3. Ray construction. A multipoint ray is constructed along the normalized converted vector that spans the full volume along the line of sight.
4. Intensity sampling. Each ray sample point within the bounds of the image reads the nearest voxel from the target image layer, which produces an intensity profile along the ray.
5. Peak selection. The maximum intensity voxel from the ray profile becomes the candidate for placement.
6. *Volume optimization (optional).* If enabled, the local 3D neighborhood of defined radius is extracted around the candidate position. The neighborhood's argmax is used as the final placement. This refinement reduces the inherent error from manual placement.

#### 1.7 Multi-round image registration and coordinate transformation

Multiple rounds of probe hybridization in sequential FISH imaging results in inter-round drift. While simpler forms of drift can be managed with global transformations, elastic nonlinear deformation provides the greatest challenge to accurately align spatial information across rounds of acquisition. zFISHer addresses this with a two-stage registration pipeline: a RANSAC-based rigid translation global estimate from nuclei centroids, followed by an optional B-spline deformable refinement on the nuclei channel intensity volumes.

**Rigid alignment via centroid RANSAC.** Rigid alignment of R1 and R2 nuclear centroid point clouds is robust to non-matching nuclei and outliers. It proceeds as follows:

1. *Vector voting coarse shift.* Pairwise difference vectors between subsampled R1 and R2 centroid sets are binned in 3D. The most-voted bin's center is selected as the coarse translation estimate.
2. *Nearest-neighbor candidate pairing.* Following application of the coarse translation shift, the nearest neighbor centroids are identified via a 3D KD-tree using scipy.spatial.cKDTree. A

search radius is used to prune implausible matches. The resulting centroid correspondence is used as input to RANSAC.
3. *RANSAC translation.* A custom 3-DOF _TranslationTransform3D model is fit with skimage.measure.ransac to prevent overfitting or implausible shear or scale from small input correspondence sizes. The final shift is mean displacement across inlier correspondences, with a reported alignment RMSD across inliers quality metric. The shift vector and RMSD is stored in the session JSON. If RANSAC fails by exceeding a deviation threshold, zFISHer falls back to the coarse vector-voting shift.
4. Canvas alignment and padding. The R1 and R2 volumes are placed in a shared canvas space with align_and_pad_images. Each round's data is padded with zeros so R1 and R2 coordinate systems share a common grid. The stored offset for each volume is used to transform puncta coordinates into the aligned space.

**Deformable B-spline refinement (optional).** This corrects subtle, nonlinear distortion that global translation cannot remove. The calculate_deformable_transform fits a B-spline transform between rigidly aligned R1 and R2 nuclei channel volumes using SimpleITK. It proceeds as follows:
1. *Downsampling.* Input volumes are downsampled by a factor of 16.
2. *B-spline parameterization.* B-splines are parameterized with a 4-node mesh in each spatial dimension.
3. *Similarity metric.* Mattes mutual information is computed with 1% voxel sampling.
4. *Optimization.* An L-BFGS-B optimizer iterates to a maximum of 100 iterations, adjusting B-spline control points, with a gradient convergence tolerance of 1e-5.
5. R2 channel warp. The fitted transform is applied to R2 channels via apply_deformable_transform. Nuclei channels use B-spline interpolation, all other channels used linear interpolation for artifact reduction, and nuclei mask layers use nearest-neighbor interpolation. Puncta point layers are not image-warped, see below.

**Outputs.** Per-channel aligned and warped volumes are saved in the aligned/ subdirectory. Checkerboard composite and deformation-field visualization layers can be generated.

#### 1.8 Consensus nuclei generation

Following registration, R1 and R2 contain independent nuclei label masks. A consensus nuclei mask is needed for colocalization analysis between channels across rounds. Nuclei centroid matching is used to associate the masks to a single ID before the masks are merged.

**Centroid matching.** R1 and R2 masks are first processed with match_nuclei_labels as follows:
1. Centroids for all masks in R1 and R2 nuclei layers are extracted with regionprops.
2. R1 centroids are used to build a 3D KD-tree. Each R2 centroid is queried for its nearest R1 neighbor. The matching threshold is adaptively computed from the distance distribution itself as median + 3*MAD (median absolute deviation).
3. R2 centroids with found partners are relabeled to match R1 IDs. Orphaned R2 IDs receive new integer labels beyond the R1 ID range. If multiple R2 centroids match the same R1, only the first keeps the shared ID. The rest received new IDs.

**Mask merging.** The new ID set is used for merge_labeled_masks, which combines masks with the same ID into a single consensus mask. Following the merge, centroids are recomputed on the final, merged masks. Two modes are provided:
- *Union.* The consensus mask contains voxels present in either round.
- *Intersection*. The consensus mask only contains voxels present in both rounds.

**Output and integration.** The consensus mask can be edited with the Mask Editor widget as described in Section 1.3. Consensus masks are saved to segmentation/Consensus_Nuclei_masks.tif and the centroid overlay is saved to segmentation/Consensus_Nuclei_masks_IDs.npy. The session and all downstream operations operate on the consensus mask and its unified ID space.

### 1.9 Puncta coordinate transformation.

Each puncta point detected prior to registration and warping is mapped into the unified canvas using transform_puncta_to_aligned_space. The transformation is different per round.

**R1 puncta layer transformation.** The canvas offset from registration padding is subtracted, placing the coordinate into the padded canvas frame.

**R2 puncta layer transformation.** Three operations are performed in sequence:

1. The rigid shift vector (from alignment) is added.
2. The canvas offset from registration padding is subtracted.
3. B-spline transform (if applicable) is applied to each R2 coordinate. The inverse is computed iteratively from the target fixed-space point, computing the residual and updating until convergence.

**Consensus nuclei reassignment.** After the coordinate transformation, the puncta are reassigned a consensus nuclei ID by indexing the consensus mask at its aligned ZYX position.

**Output and integration.** The transformed puncta layers' CSV files are written into the reports/ subdirectory with coordinates in the unified canvas space and IDs from the assigned consensus nuclei. These are used as input for colocalization analysis.

### 1.10 Colocalization analysis

zFISHer defines colocalization as the co-occurrence of two (pairwise) or three (tri-channel colocalization) from different channels within a defined physical distance (micrometers). The operation is performed in the aligned canvas space following registration and warping.

**World-coordinated conversion.** All distance calculations operate on world coordinates and not voxel indices to remain consistent across datasets with different voxel sizes. Each point's pixel position is converted to physical space (world = data · scale + translate). Scale contains the voxel dimensions (µm) from OME-XML metadata and translate accounts for canvas offset introduced during registration padding.

**Pairwise colocalization.** For each defined pair, symmetric radius search is performed using scipy.spatial.cKDTree.query_ball_tree. All target points within a threshold distance in relation to the source point are identified. Pairs are deduplicated via a canonical ordering on layer names and indices so A→ B and B→A queries are not counted twice for the same physical relationship event. Note that a point can exist in multiple pairwise hits, as long as it is paired with a different point not already identified as a positive colocalization event. Each recorded pair exists as a one-row-per-hit table (both layer names and point indices, consensus nucleus ID(s), 3D coordinates, computed distance, threshold distance) suitable for downstream metric generation.

**Tri-channel colocalization.** zFISHer calculates tri-channel colocalization using a greedy matching algorithm. An anchor channel is defined alongside two comparison channels (A and B). A single distance threshold is applied for both anchor→A and anchor→B. This approach is intended to represent a biological model, where factors associate to a single spatial locus. The algorithm proceeds as follows:

1. *Anchor lookup tree.* A KD-tree is built on anchor-channel world coordinates.
2. *Per-channel nearest-anchor assignment.* Every channel A and channel B point is assigned to its closest anchor point. Assignments above the threshold are discarded.
3. *Candidate enumeration.* For each anchor with at least one A and B point assigned, all (A,B) combinations are listed exhaustively.
4. *Greedy triplet selection.* Candidates are sorted by total distance cost function (chA_dist + chB_dist). The triplet with the lowest total is accepted, and the consumed channel A and B puncta are marked as used. This repeats until the pool has no remaining valid triplets.

Each recorded tri-channel colocalization event exists as a one-row-per-hit table (layer names, point indices, consensus nucleus ID(s), 3D coordinates, computed distance, threshold distance) suitable for downstream metric generation.

**Comprehensive point distance generation.** A separate calculate_distances routine calculates the distance of every point in every layer to its nearest neighbor in every other layer, independent of any colocalization rule. This produces a comprehensive background distribution for reporting nearest-neighbor distances and additional metrics and aggregates.

#### 1.11 Export and visualization

The output of zFISHer is intended to export a comprehensive report of the analysis into easily parsed data and metrics for additional analysis, in addition to publication-ready elements such as screenshots.

**Report export.** The comprehensive report generated by export_report is written to the reports/ subdirectory as a single multi-sheet workbook. The sheets are:

- *Distances.* The full nearest-neighbor table of all point-to-point distances across all layers.
- *Colocalization.* Pairwise colocalization hits. One row per pair.
- *Tri-channel colocalization.* Tri-channel colocalization hits. One row per triplet.
- *ROI per nuclei.* Per-consensus-nucleus puncta counts, aggregated across channels.
- *Stats.* Per-channel summary statistics including count, mean distance, standard deviation, and coefficient of variation.
- *Distribution.* Binned distance distributions.
- *Parameters.* The full rule set and session metadata. This provides a complete analysis provenance record.

**Scale bar.** A custom DraggableScaleBar overlay renders a physical-unit scale bar whose length is computed from the current layer's voxel spacing in micrometers and the current canvas zoom. The pixel-length relation to the physical unit can also be displayed. The bar can be repositioned for image capture.

**Arrow annotation overlay.** A transparent ArrowOverlay overlay allows users to draw arrows directly on a specific slice of the canvas in 2D mode. The arrows are stored in data coordinates rather than screen pixels, allowing them to directly translate to proper 3D space in 3D viewing mode.

**Screenshot capture.** Screenshots of the current view can be captured with a one-click export to PNG and include scale bar and arrows if applicable. Captures are saved to the captures/ subdirectory with incrementing filenames.  Additionally, a configurable hotkey allows a region-selection mode to export a crop region of the canvas.

### **Supplementary Methods 2.** Sequential XPO1 IF & HOXB DNA FISH Sample Preparation and Imaging.

**Probe preparation.** Green labeled HOXB (Empire Genomics Corp / RPC1-11 29G13 / 17q21 / chr17:46602703-46770717 Hg19) was used as the DNA FISH probe. XPO1 Rabbit mAb (CST #46249S) was used as the primary antibody at a dilution of 1:100 and AF647 goat, anti-rabbit (Invitrogen #A21245) was used as the secondary antibody at a 1:200 dilution.

**Sample processing.** Single cells from cell line Molm13 cells were applied to glass slides by cytocentrifugation. Slides were fixed in 1% PFA in PBS for 5 minutes followed by 1% PFA in PBS plus 0.05% Igepal for 5 minutes, and stored in 70% ETOH at -20°C. Slides were rinsed in PBS and blocked in 200µl blocking buffer (2X SSC and 1% BSA) and then incubated with the XPO1 Ab (1:100 dilution) in antibody diluent (2X SSC and 1% BSA) for 45 minutes at RT in the dark. Slides were then washed in PBS and incubated with the AF647 Ab (1:200) in antibody diluent for 45 minutes at RT in the dark. Slides were then washed in PBS and stained with Vectashield containing DAPI and imaged for IF (1-17-24Adecon.nd2). Slides were then washed in PBS and digested with RNaseA for 45 minutes at 37°C. Cross linking with 4% PFA for 9 minutes was then performed, followed by treatment in 0.2N HCL for 10 minutes on ice. Slides were then denatured in 70% formamide, 2X SSC at 80°C for 8 minutes followed by ETOH dehydration series. Denatured HOXB probe mixed with hybridization buffer (Empire Genomics Corp) was applied to the slides and hybridized at 37°C overnight. Slides were then washed in 50% formamide, 2X SSC at 37°C for 5 minutes and stained with Vectashield containing DAPI and imaged again using the same coordinates as had been imaged in the first set of IF images (1-19-24Fdecon.nd2).

**Image capture.** Imaging was performed in 3D using a widefield fluorescence microscope (Nikon Eclipse E8000) equipped with Nikon Nis Elements version 4.2 software and Autoquant 3D deconvolution, a Lumencor Sola light engine, and a Sutter Lambda 10-3 filter wheel controller. Images were deconvolved using 3D Automatic with number of rounds set to automatic. Images were acquired using 0.15-micron plane spacing and a 60X planapochromatic objective with a numerical aperture of 1.4. The number of Z steps for 1-17-24Adecon.nd2 was 60 with deconvolution iterations of 58, 90, 90; exposures were 300ms (DAPI), 300ms (FITC), and 1s (CY5). The number of Z steps for 1-19-24Fdecon.nd2 was 71 with deconvolution iterations of 73, 38, 90; exposures were 300ms (DAPI), 500ms (FITC), 1s (Cy5). Raw image files (1-17-24Adecon.nd2 for the XPO1 IF and 1-19-24Fdecon.nd2 for the HOXB DNA FISH acquisitions) are archived on Zenodo at https://doi.org/10.5281/zenodo.20288536.

### **Supplementary Methods 3.** zFISHer Analysis Workflow Applied to Sequential XPO1 IF & HOXB DNA FISH Example Dataset.

The example analysis presented in Application and Supplementary Methods 2 was performed using zFISHer's interactive environment with the parameters and manual curation described below.

**Input imports.** The analysis pair was imported as Round 1 (HOXB DNA FISH, 1-19-24Fdecon.nd2; 3 channels, 71 z-slices) and Round 2 (XPO1 immunofluorescence; 1-17-24Adecon.nd2; 3 channels, 60 z-slices). The files were imported as ND2 and automatically converted to OME-TIFF. The parsed OME-XML metadata yielded voxel dimensions of dz = 0.15 um, dy = dx = 0.108 μm with an anisotropy ratio (dz/dx) of 1.38 for downstream correction.

**Nuclear segmentation.** The modified watershed algorithm was applied to each round's DAPI channel independently for nuclear segmentation, using the default zFISHer parameter configuration. Segmentation produced 55 reference nuclei in the consensus mask after mask editor cleanup.

**Puncta detection.** Automated spot detection was performed on the unaligned channels using the Local Maxima algorithm with the parameter configurations described as follows. The XPO1 (Cy5) channel was set to a sensitivity threshold of 0.03 and the HOXB (FITC) channel was set to a sensitivity threshold of 0.07. All other detection parameters were left at the default zFISHer configuration. White top-hat background subtraction was not used. Detection was constrained to within the relevant DAPI channel mask to exclude extranuclear puncta. Puncta located within five frames of the image stack boundaries were excluded prior to analysis, as these were determined to be artifacts from centrifugation during slide preparation. Specifically, spun-down protein and probe material present on the slide surface, external to the nucleus, were removed. The Fishing Hook tool was used to add a single point in the FITC channel, and no manual curation was performed on the Cy5 channel.

**Alignment and warping.** Cross-round alignment was performed using zFISHer's two-stage registration pipeline: RANSAC-based rigid translation on per-round nuclear centroids (default configuration parameters), followed by B-spline elastic warping on the R2 nuclear channel, linear interpolation on the remaining R2 image channels, and inverse-warp coordinate transformation on the R2 puncta coordinate layers.

**Consensus nuclei generation.** The warped and aligned R1 and R2 nuclear DAPI masks were aligned into a consensus mask using the Intersection mode (default configuration parameters). A total of 11 nuclei were deleted from the consensus nuclei masks using the manual hover-delete mode, which were primarily nuclei that were not fully captured along the image boundaries. Puncta coordinates outside the consensus nuclei of all channels were filtered out using the puncta cleanup tool.

**Colocalization analysis.** Pairwise colocalization between the FITC (HOXB) and Cy5 (XPO1) channels was performed with a 1.0 µm distance threshold. This yielded 112 FITC-Cy5 nearest neighbor possible pairs. Of the possible pairs, 11 pairs were found to be below the threshold of 1 μm, distributed among 8 unique FITC and 11 Cy5 partners. The 8 FITC positive hits were counted as colocalized.

**Report generation.** The final analysis output, session, and report containing the per-hit pairwise tables, per-nucleus aggregates, summary statistics, and parameter set, was exported to a single multi-sheet report. The results are consistent with the expected biological outcome (Application; Figure 1E and 1F).
